# ThruTracker: Open-Source Software for 2-D and 3-D Animal Video Tracking

**DOI:** 10.1101/2021.05.12.443854

**Authors:** Aaron J. Corcoran, Michael R. Schirmacher, Eric Black, Tyson L. Hedrick

**Affiliations:** Department of Biology, 1420 Austin Bluffs Blvd, University of Colorado, Colorado Springs, Colorado, USA; Bat Conservation International, 500 N Capital of TX Hwy. Bldg. 1. Austin, Texas, USA; Canebrake Environmental Services, LLC. 7405 New Forest Lane, Wake Forest, NC 27587; Department of Biology, University of North Carolina, 120 South Road, Chapel Hill, North Carolina, USA

**Keywords:** Animal Flight, Animal Movement, Movement Ecology, Population Monitoring, Flight Biomechanics, Video Detection

## Abstract

1. Tracking animal movement patterns using videography is an important tool in many biological disciplines ranging from biomechanics to conservation. Reduced costs of technology such as thermal videography and unmanned aerial vehicles has made video-based animal tracking more accessible, however existing software for processing acquired video limits the application of these methods.
2. Here, we present a novel software program for high-throughput 2-D and 3-D animal tracking. ThruTracker provides tools to allow video tracking under a variety of conditions with minimal technical expertise or coding background and without the need for paid licenses. Notable capabilities include calibrating the intrinsic properties of thermal cameras; tracking and counting hundreds of animals at a time; and the ability to make 3-D calibrations without dedicated calibration objects. Automated 2-D and 3-D workflows are integrated to allow for analysis of largescale datasets.
3. We tested ThruTracker with two case studies. The 2-D workflow is demonstrated by counting bats emerging from bridges and caves using thermal Videography. Tests show that ThruTracker has a similar accuracy compared to humans under a variety of conditions. The 3-D workflow is shown for making accurate calibrations for tracking bats and birds at wind turbines using only the wind turbine itself as a calibration object.
4. ThruTracker is a robust software program for tracking moving animals in 2-D and 3-D. Major applications include counting animals such as bats, birds, and fish that form large aggregations, and documenting movement trajectories over medium spatial scales (∼100,000 m^3^). When combined with emerging technologies, we expect videographic techniques to continue to see widespread adoption for an increasing range of biological applications.

## 1. Introduction

Video-based animal tracking is a widely used tool in fields as diverse as biomechanics, animal behavior, ecology, and population monitoring (Dell et al., 2014). The reduced price of thermal video cameras, high-speed cameras, and other technologies such as unmanned aerial vehicles (UAVs) has expanded dramatically the capabilities of investigators studying animal movement in the lab and in the field (Jackson, Evangelista, Ray, & Hedrick, 2016). This has placed great demand for video processing software.

Various video tracking tools are available for two-dimensional (2-D), and three-dimensional (3-D) animal tracking. Two-dimensional tracking is often used for studies of one or multiple individuals under controlled laboratory settings. Two-dimensional tracking can also be useful for surveying animal populations such as bats or birds and for monitoring their migratory behavior (Kunz et al., 2009). Algorithms differ in how they detect animals and how they connect detections between frames to form spatiotemporal tracks. For example, some software uses background subtraction for detection (Rodriguez et al., 2018) while other programs use adaptive thresholding (Sridhar et al., 2019). One popular program uses image recognition to help track individuals (Pérez-Escudero et al. 2014). Deep learning based approaches have recently become popular, particularly for marker-less tracking of animal body parts (Mathis et al., 2018; Pereira et al., 2019).

Three-dimensional animal tracking uses two or more cameras to triangulate animal positions. This requires synchronized video acquisition, careful calibration of the optical properties of cameras [i.e.2camera extrinsics). Several software programs are available for generating 3-D calibrations and tracking animals in 3-D (Noldus et al., 2001; Hedrick, 2008; Theriault et al., 2014; Jackson et al., 2016; Knorlein et al., 2016; Nath et al., 2019).

Originally, 3-D calibrations required that an object with several markers at known 3-D positions be placed within the field of view of each camera (Abdel-Aziz & Karara, 1971). More recent workflows produce calibrations by moving an object with a recognizable 2-D pattern [“checkerboard calibration” (Zhang, 2000)] or with two markers at a fixed distance (“wand calibration”) through the calibration volume (Theriault et al., 2014). This allows one to set the scale of the scene using the wand length or pattern scale and to estimate the accuracy of the calibration based on the variation of reconstructed wand lengths. However, researchers are increasingly interested in studying animals within large volumes in the natural environment where it is difficult to deploy calibration objects or where doing so might disturb the animals under study (Evangelista, Ray, Raja, & Hedrick, 2017; Corcoran & Hedrick, 2019). In addition, much of the currently available 3-D tracking software requires manual digitization of objects in the videos, which limits the amount of data that can be processed. Finally, calibration procedures are not widely available for determining the intrinsic properties of thermal cameras, preventing their widespread use for 3-D tracking applications.

Here we present ThruTracker, a free and open-source software package for high-throughput 2-D and 3-D animal tracking. ThruTracker provides an app-based environment (i.e., no coding required) with all the tools necessary to track animals under a variety of conditions with light-based or thermal cameras. ThruTracker is coded in MATLAB v2020b (Natick, MA, USA) and compiled versions of the software are provided so that it can be installed and run for free without any licenses. We provide source code under a 3-clause BSD license for those who may want to customize the software for their own purposes. ThruTracker is also compatible with any tracking software that outputs 2-D coordinates, and it exports data in standard text formats that can be imported into other software for further processing or statistical analysis.

We first provide an overview of the 2-D and 3-D track generation process, before demonstrating the use of ThruTracker with two test cases—tracking bats at wind turbines and counting bat emergences from bridges and caves using a thermal camera flown with an Unmanned Aerial Vehicle (UAV) or recording from the ground. We conclude by discussing applications of 3-D tracking and current limitations. A step-by-step manual for ThruTracker along with additional recommendations for camera setups and use of the software is available on our website (www.sonarjamming.com/thrutracker).

## 2. Materials and Methods

### 2.1 Workflow for 2-D and 3-D Tracking

ThruTracker can be used either for 2-D or 3-D video tracking. Two-dimensional tracking does not require that the videos be calibrated and is therefore a much simpler procedure. Three-dimensional tracking requires calibration of the camera properties such as focal length and lens distortion (intrinsic calibration) and determination of the positions and orientations of the cameras for each scene where they are deployed (extrinsic calibration). The workflow for 3-D track generation is shown in Figure 1.

**Figure 1.**
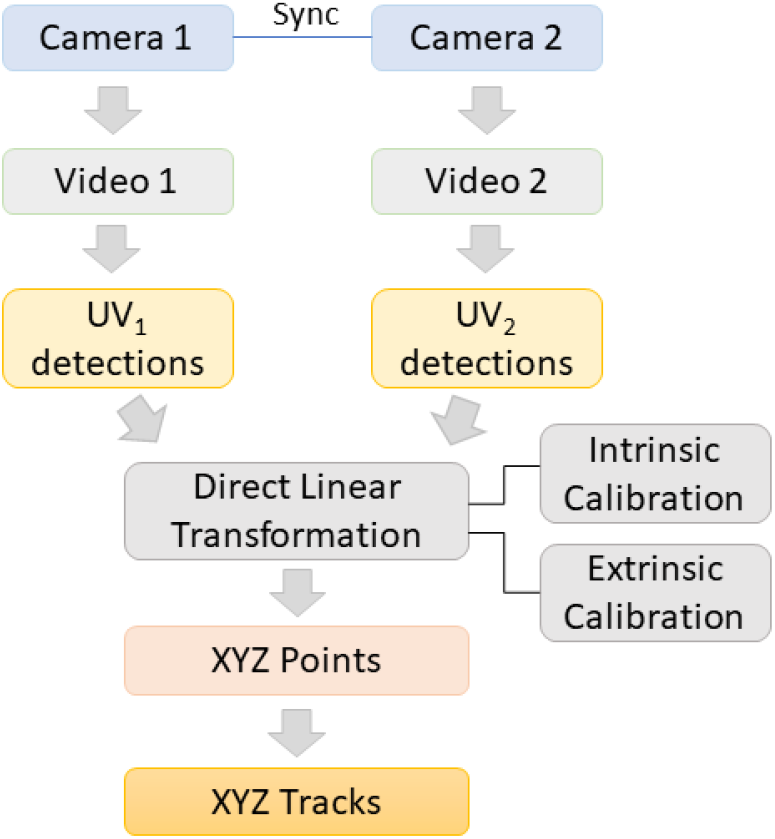
Overview of 3-D track generation procedure. Two or more cameras acquire synchronized video, which are each processed to generate 2-D (UV) detections. Intrinsic and extrinsic camera calibrations are used to generate direct linear transformation (DLT) equations that transform UV coordinates into real-world 3-D or XYZ points for each detection. XYZ points are then stitched together across frames to generate 3-D tracks.

#### Step 1: Video Acquisition

Synchronized videos are acquired using two or more cameras. Some cameras, such as FLIR A65 thermal video cameras (FLIR Systems, Inc., Wilsonville, OR) used in our testing, have dedicated electronic inputs that allow digital signals to synchronize the shutters of each camera. An alternative for cameras without sync ports is to use audio signals that are broadcast to each camera (Jackson et al., 2016). Audio synchronization of cameras does not allow for shutter synchronization, therefore there will be a time offset of up to half the frame rate even after synchronization. Recording at 30 Hz, a half-frame synchronization accuracy has been sufficient to achieve highly accurate calibrations for animals moving at low to moderate pixel speeds (Corcoran & Hedrick, 2019). Audio synchronization may also require fine-tuning since some camera models may not keep the audio and video outputs in perfect alignment.

#### Step 2: Object detection

After videos are acquired, videos from each camera are processed to detect moving objects in each frame. ThruTracker uses a Gaussian mixture-based background subtraction algorithm implemented in OpenCV [“BackgroundSubtractorMOG2”, (Zivkovic, 2004; Zivkovic & Van Der Heijden, 2006)]. A blob detector is then used to isolate detections and the blob centroids are used as detection coordinates. A simplified interface allows the user to rapidly select and modify detection settings for their application. Settings include: 1) selecting which frames to process, 2) minimum and maximum object size in pixels, 3) sensitivity for adjusting the threshold for discriminating foreground from background, 4) Number of background frames used for generating the rolling background image, and 5) target object diameter. This last option determines the size of a 2-D gaussian filter that is applied to the image, which helps reduce noise and isolate closely spaced animals. Each frame of the video is processed for detections before detections are linked together into 2-D or 3-D tracks in the proceeding steps. Two-dimensional tracking applications can skip to step 6.

#### Step 3: Intrinsic camera calibration

For 3-D tracking applications, one must determine the camera’s intrinsic properties, including the focal length, principal point and lens distortion (Abdel-Aziz & Karara, 1971; Hartley & Zisserman, 2004; Lourakis & Argyros, 2009). This information can be used to map any pixel in the camera image into a vector in a camera-based frame. Typically, this calibration can be done once per camera and lens combination in a laboratory and the values should be similar between cameras and lens of the same make and model. In some special circumstances one might need to calibrate each individual camera-lens pair for increased accuracy. Cameras with variable focal length (i.e., a zoom lens) typically need to be calibrated separately at each focal length used for recording.

We use MATLAB’s built-in camera calibratio functions (Bouguet, 1999) with some modifications to calibrate thermal images (Yahyanejad, Misiorny, & Rinner, 2011). We recommend using MATLAB’s built-in camera calibrator app or freely available functions in OpenCV or Argus (Jackson et al., 2016) for calibrating light-based cameras.

Traditionally, calibration procedures for light-based cameras rely on detecting the corners of a checkerboard pattern. With thermal imaging, one must first heat the checkerboard so that the darker squares will be hotter than the lighter squares because of higher light absorption (Figure 2a). For example, we achieved this by moving two 100-watt lamps over the checkerboard pattern for about 20 seconds before taking thermal images.

**Figure 2.**
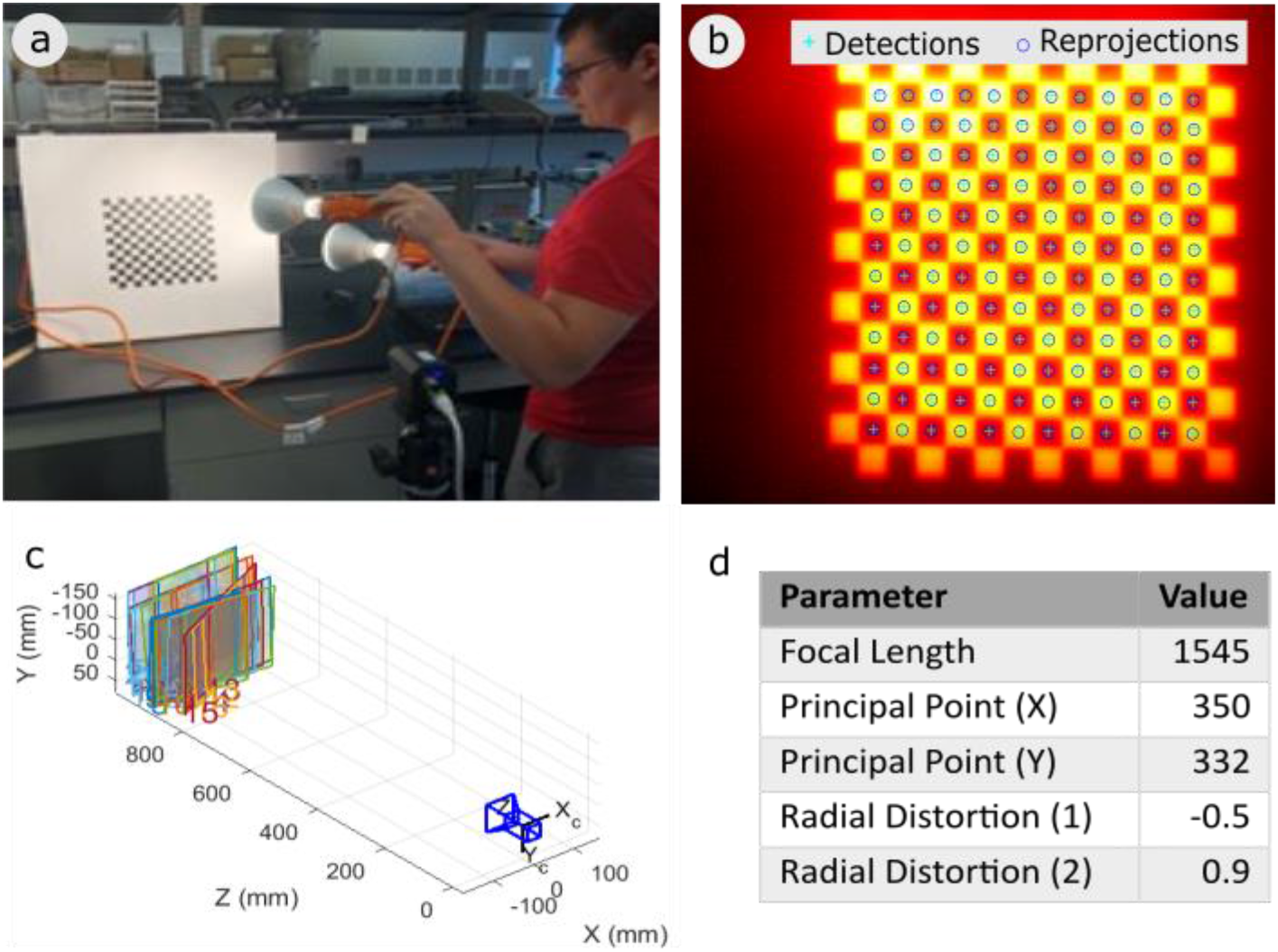
Thermal camera intrinsic calibration. (a) An experimenter heats a checkerboard pattern with two 100-watt lamps. (b) Example thermal image of a checkerboard pattern with detected and reprojected square centers. (c) Graphical depiction of the camera and 28 images of the checkerboard pattern taken at different positions and orientations. (d) Example output of intrinsic camera parameters for a Flir A65 camera.

However, uneven heating combined with rapid conduction of heat through the checkerboard image increases the difficulty of detecting the corners between light and dark squares (or in this case between cold and hot squares; Figure 2b). Instead of detecting the corners, the cooler portions of the image are dilated so that each hot square is reduced in size and no longer connected to adjacent hot squares. A blob-detector is then used to detect the centers of the hot squares. The pixel values of the image are then inverted, and the procedure is repeated to detect the centers of the cold squares. This is repeated for 20-30 images taken at different positions and orientations relative to the camera. The resulting data are used by the software to calculate the camera’s intrinsic parameters (Figure 2c, d).

#### Step 4: Extrinsic camera calibration

A second calibration procedure is required for generating 3-D tracks for each recording setup—that is, any time the cameras are moved even slightly. The extrinsic calibration determines the positions of the cameras in real-world 3-D space and their orientations [roll, yaw and pitch; (Abdel-Aziz & Karara, 1971)]. Together with the intrinsic calibration data, the extrinsic calibration allows one to map objects that are detected in two or more cameras to real-world 3-D coordinates.

As noted previously, there are several approaches for generating extrinsic calibrations. ThruTracker has two options: a wand-based procedure that is based on methods used for easyWand (Lourakis & Argyros, 2009; Theriault et al., 2014) and a procedure that is based exclusively on background points. One can also import calibrations made using DLTdv or easyWand (Hedrick, 2008; Theriault et al., 2014). Background points include any object that is visible within both cameras such as tips of tree branches, wind turbines, or animals. Moving objects can be used if the images are taken synchronously between cameras. This background point procedure [also known as structure from motion (Schönberger & Frahm, 2016)] means that 3-D calibrations can be made without dedicated calibration objects within the field of view. In effect, any object visible by two or more cameras can be used as a calibration object. Two additional components of the calibration must be determined when no dedicated calibration objects are used: the scale of the scene and the gravitational axis. The scale can be set by specifying two points in the scene that are at a known distance from one another. Alternatively, one can use the distance between cameras to set the camera scale.

Second, the gravitational axis can be set by measuring the inclination angle of one of the cameras using an inclinometer. This value is input into ThruTracker’s calibration app during the calibration procedure. With these options, one can obtain calibration data rapidly in the field with only a few measurements and with no requirement for deploying calibration objects. This is especially helpful for situations where it is not feasible to deploy calibration objects or where their use would disturb animals under study.

A notable down-side of not using calibration objects is the absence of objects at known positions that can be used to check the accuracy of the calibration. Therefore, it is important to conduct tests of the calibration procedure using objects at known distances. For example, in our test using wind turbines below, we measured the variation in the reconstructed lengths of the turbine blades at different times and positions throughout our recording. Another alternative would be to conduct test setups in the field at locations where it is easier to deploy calibration objects such as a wand.

#### Step 5: Generating 3-D Points

Three-dimensional points are generated after objects have been detected in videos from each camera and intrinsic and extrinsic calibrations are completed. For each set of synchronized video frames, theoretical 3-D points are created from all combinations of 2-D detections across cameras. Each putative 3-D point has an associated direct linear transformation (DLT) residual. The DLT residual is the distance in pixels between the observed image coordinates of a marker and the “ideal” image coordinates computed from the estimated 3-D location of the marker and the calibration information for the camera that captured the image. If we visualize each 2-D detection as a vector in 3-D space with its origin at the camera, the residuals indicate how closely a given set of 2-D detections and their associated vectors come to crossing in 3-D space. The algorithm starts by creating 3-D points based on sets of 2-D points with the lowest residuals and removing those points from the available pool. It proceeds until there are no more 3-D points with residuals below the specified threshold.

#### Step 6: Generating 2-D and 3-D Tracks

Two-dimensional and Three-dimensional points are stitched together across frames to make tracks. Each 2-D or 3-D point in the first frame is a putative track. Detections in each proceeding frame are assigned to existing tracks, or if no assignment is made, they become the beginning of a new track. A Kalman filter is applied to each track to predict the position of the track in the next frame. The distance between each detection and the predicted positions are computed. This is done both in 2-D and in 3-D to calculate a cost matrix. A Hungarian algorithm is used to determine the combination of assignments that minimizes cost across the assignment matrix (Kuhn, 1955). Finally, a threshold is specified in ThruTracker such that assignments are only made if their cost is below the threshold. One can specify the number of frames between detections that are allowed (i.e., gap distance) before a track is terminated. One also specifies the minimum number of detections required for a track to be retained. Longer gaps and smaller minimum numbers of detections increase the number of tracks that are retained but increases the number of false positive tracks generated from noise.

#### Step 7: Data visualization, analysis and classification

ThruTracker offers multiple tools for visualizing and processing tracks. One can rapidly toggle between tracks and use shortcut keys to classify them into different categories. For example, for wind turbine applications, the tracks can be labeled “bird”, “bat”, “airplane”, “noise”, etc. Another option allows all the tracks to be visualized at once. Tracks can then be selected as groups and classified based on their positions, start or end points. This tool is helpful for selecting tracks based on their location, as exhibited in the bat emergence case study presented below. Another option allows the user to draw a rectangle over the camera image to count exits and re-entries as animals pass into or out of the rectangle. This is useful, for example, when counting bats exiting a cave roost. Resulting 2-D, 3-D and track data can be exported into CSV files for use in other analysis programs.

### 2.2 Case Studies

#### 2.2.1 Case Study 1: Counting bat exits from bridges and caves using thermal imaging

We used ThruTracker to count bats emerging from bridges using a DJI Zenmuse XT2 thermal camera with a 13 mm lens (45-degree field of view) suspended from a DJI Matrice 300 drone (SZ DJI Technology Co., Shenzhen, China). Because this analysis was done in 2-D there was no need for intrinsic or extrinsic calibrations. The drone was flown at altitudes of 50 m and 80 m above a bridge known to be a roost location for big brown bats (*Eptesicus fuscus*) in August of 2020 near Burnsville, NC, USA. We also counted gray bats leaving caves using thermal cameras placed at ground level. Videos of gray bats were provided by the US Fish and Wildlife Service.

Our goals were 1) to determine the maximum distance at which bats could be detected and 2) to compare manual counts of emergences with those produced using ThruTracker. The bridge recordings provide a test of relatively low numbers of bats counted near the limits of their detection range. The cave recordings test detection of large numbers with high rates of occlusion.

Videos were processed in ThruTracker with the following parameters: sensitivity, 35; background frames, 20; Min object pixels, 1; max object pixels 100; min track length 5, max gap length 5; match threshold 10. After detections were made in ThruTracker, the applet TrackSelector was used to rapidly select tracks that originated near the edge of the bridge. Manual observers used VirtualDub software to play videos at 50% of normal speed and paused and reviewed videos frame by frame, as necessary.

We processed two videos taken at heights of 50 m and 80 m above the bridge and two videos representing different bat densities at caves (Table 1). Videos were not meant to census the entire emergences, but rather provide data for comparing detection abilities.

**Table 1.**
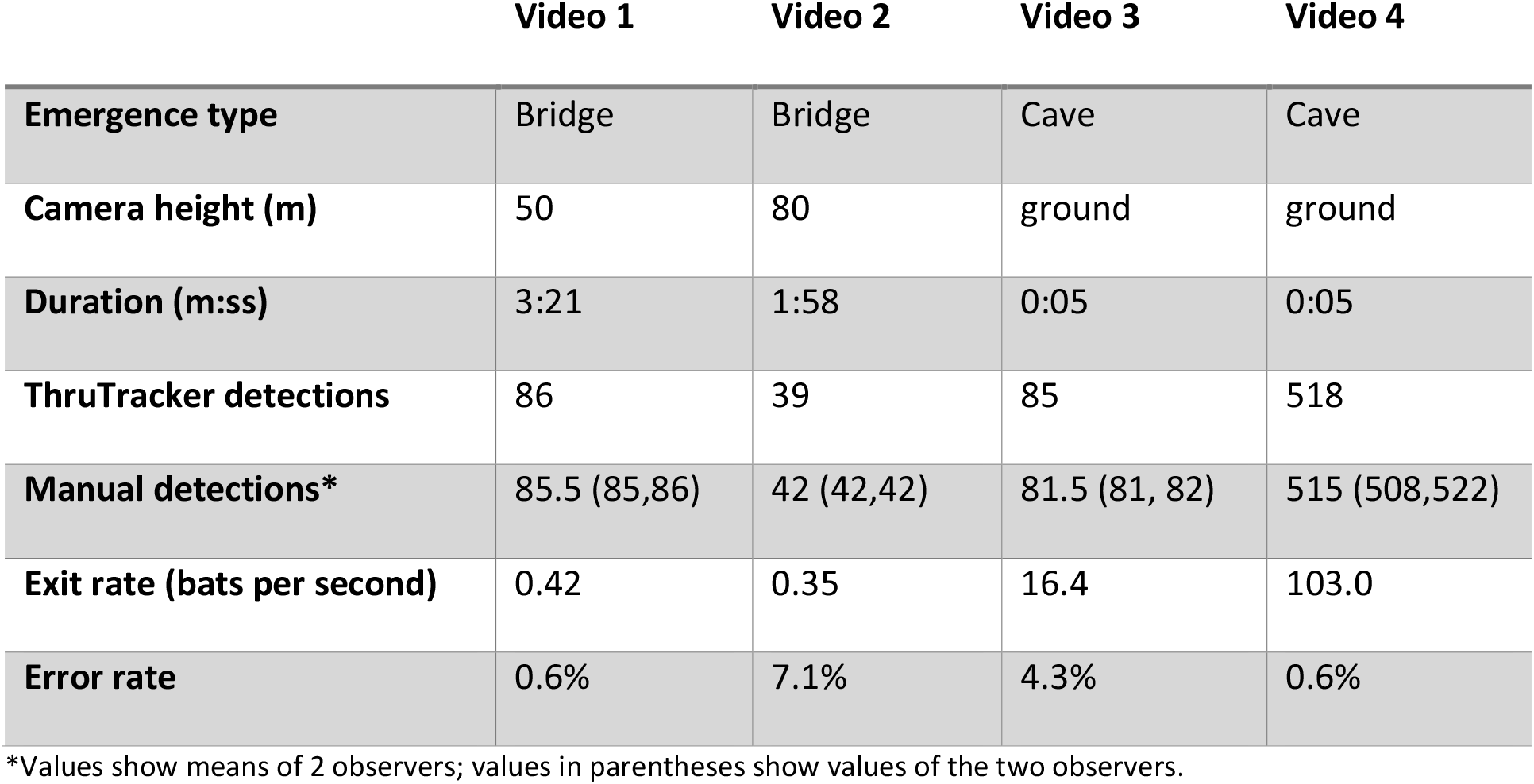
Comparison of manual and automated bat counts using thermal videography.

|  | Video 1 | Video 2 | Video 3 | Video 4 |
| --- | --- | --- | --- | --- |
| **Emergence type** | Bridge | Bridge | Cave | Cave |
| **Camera height (m)** | 50 | 80 | ground | ground |
| **Duration (m:ss)** | 3:21 | 1:58 | 0:05 | 0:05 |
| **ThruTracker detections** | 86 | 39 | 85 | 518 |
| **Manual detections\*** | 85.5 (85,86) | 42 (42,42) | 81.5 (81, 82) | 515 (508,522) |
| **Exit rate (bats per second)** | 0.42 | 0.35 | 16.4 | 103.0 |
| **Error rate** | 0.6% | 7.1% | 4.3% | 0.6% |

\*Values show means of 2 observers; values in parentheses show values of the two observers.

#### 2.2.2 Case Study 2

We tested ThruTracker’s ability to make 3-D calibrations for tracking 3-D flights of bats and birds at wind turbines. Studying animal movements at wind turbines is a problem of considerable interest, especially for bats who are being killed in large numbers for mostly unknown reasons (Arnett & Baerwald, 2013). Thermal imaging has been used for studying bats at wind turbines in 2-D (Horn, Arnett, & Kunz, 2008; Cryan et al., 2014) and 3-D (Kinzie et al., 2018; Schirmacher, 2020). Proprietary software for 3-D tracking of bats at wind turbines (was also recently published (Matzner, Warfel, & Hull, 2020). Here we report on our field camera setup, intrinsic and extrinsic calibration. Because we conducted these tests under cold winter conditions with few bats or birds present, we did not obtain 3-D animal tracks. For example tracks using a similar approach, see Schirmacher, 2020 (Schirmacher, 2020).

We tested the calibration setup using two Flir A65 thermal cameras with 25-degree lenses synchronized with electronic inputs at an experimental wind turbine at the National Renewable Energy Laboratory (NREL) in Golden, CO during December 2019. The cameras were placed 33 m apart from one another and 40 m from the base of the turbine’s monopole. Cameras were aimed slightly below the wind turbine’s nacelle, which was 80 m off the ground. This resulted in an inclination angle of 62.2 degrees for our reference camera.

## 3. Results

### 3.1 Case Study 1: Counting bat exits from bridges and caves using thermal imaging

ThruTracker detected similar numbers of bats compared to the manual observers in all test cases, with error rates of 0.6-7.1% (Table 1; Figure 1; Supplemental Video 1. Thermal video taken from a UAV showed that the 50 m recording height was comfortably within the detection range for the bats. Bat detections had a size of 5.2 ± 3.4 pixels (mean ± st. dev.) and tracks extended across most of the image (Figure 3a) including over water and over the bridge. However, bats failed to be tracked over some land areas where they lacked contrast with the background (e.g., see tracks terminating as they approach land area in bottom right portion of Figure 3a). Manual inspection of videos at these locations found that bats were not readily visible to the human eye, so this appears to be a limitation of the thermal imaging, not the detection algorithm.

**Figure 3.**
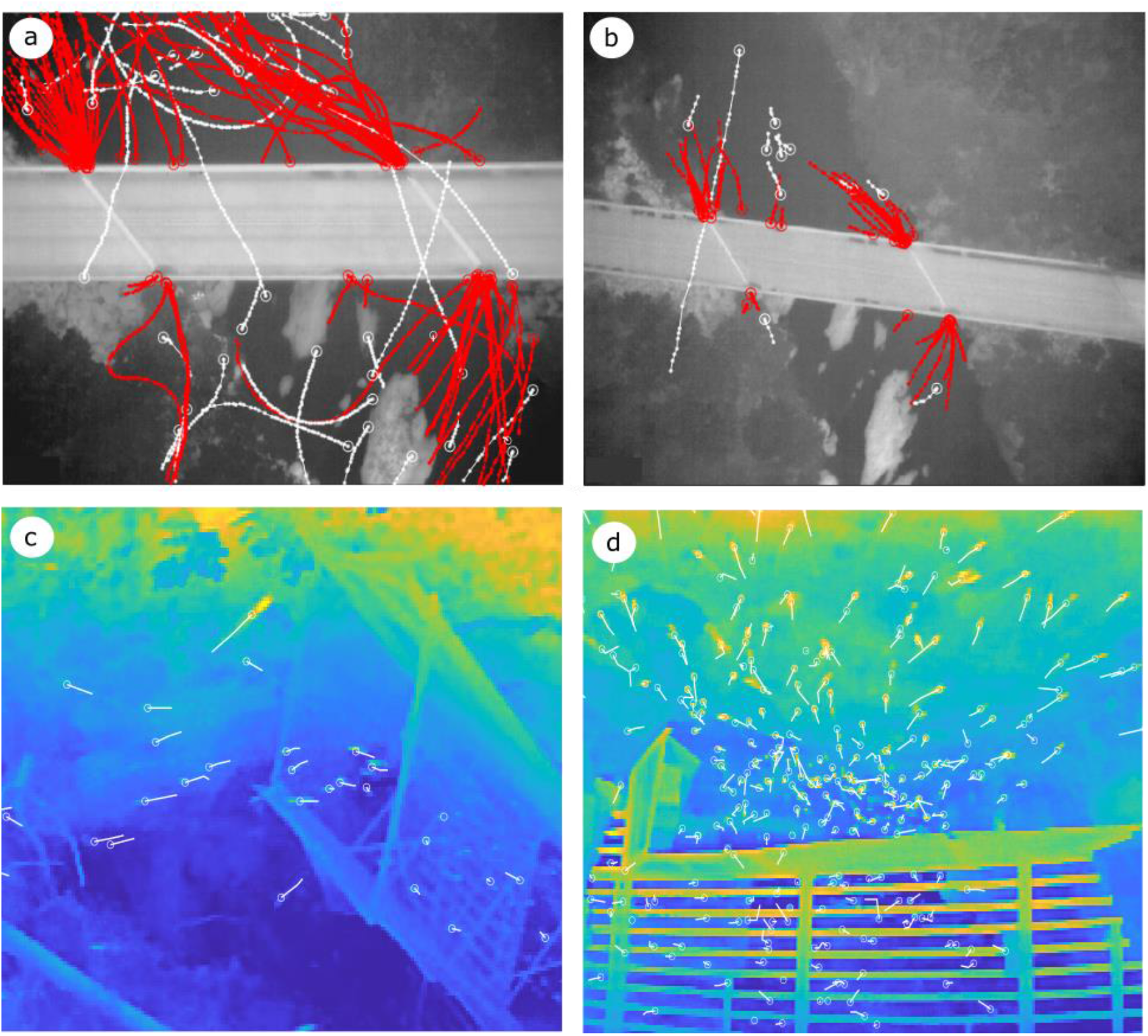
Example ThruTracker detections of bats leaving a bridge (a, b) and cave (c, d). Big brown bats (*Eptesicus fuscus*) were filmed exiting a bridge using a thermal camera on a UAV flying at 50 m (a) and 80 m altitude (b). Red tracks indicate exits and white tracks indicate other detections. In (a, b) circles indicate the starting point of tracks to highlight departures from the bridge and the entirety of all tracks are shown. In (c, d) detections from a single frame are shown (circles) with tracks indicating movement over the two previous frames. See table 1 for statistics.

The 80 m recording height was much closer to the detection limit for *E. fuscus* bats using this recording setup. Bats had a detection size of 1.9 ± 1.1 pixels. Visual inspection of videos revealed that bats were only barely visible and that they were not detectable to the human eye within a short distance of their emergence. This is reflected in the ThruTracker tracks that terminate over open water some distance from the bridge exits (Figure 3b). This may have resulted from bats dropping to a lower altitude (and further distance from the camera) as they left the bridge, as was visible in other camera views taken at an oblique angle relative to the ground. ThruTracker performed slightly worse under these at the 80 m height than the 50 m height, but still detected 93% of tracks that were detected manually (Table 1).

Counts of gray bats (*Myotis grisescens*) exiting caves were used to test of ThruTracker’s multi-object tracking abilities. ThruTracker achieved error rates of 4.3% and 0.6% for the two test videos, which we estimated to have exit rates of 16.4 and 103.0 bats per second (Table 1; Figure 3c, d; Supplemenatal Video 2).

### 3.2 Case Study 2: Calibrating large 3-D volumes using wind turbines

In our second case study, we demonstrate ThruTracker’s 3-D workflow for calibrating large spatial volumes using only the turbine itself as a calibration object. We calibrated the FLIR A65 thermal cameras (640 x 512 pixel resolution) using the intrinsic calibration method described above. Example images and resulting intrinsic camera parameters are shown in Figure 2. We used 28 checkerboard images, with an average reprojection residual error of 0.69 pixels (range 0.48-0.83 pixels).

We generated an extrinsic calibration in ThruTracker using 67 points from the wind turbine as background points (Figure 3). These included hot spots, corners and an anemometer on the nacelle, and turbine blade tips. Points were digitized manually using DLTdv8 (Hedrick, 2008). Efforts were made to select points that were 1) clearly visibly in both cameras, 2) distinct points in 3-D space, such as small hot spots or sharp edges of objects, and 3) covering a broad range of 2-D and 3-D positions. We excluded six points because they had DLT residuals > 3 pixels. The remaining calibration had mean reprojection errors of 0.63 pixels.

We set the scale of the scene using the distance between the two cameras (33 m) and the gravitational axis was set using the inclination angle of the second camera (62.2 degrees). The resulting calibration had a volume of 235,597 m3 assuming a maximum detection distance of 200 m. The maximum detection range is likely less than 200 m for small bats (<30 g) with this camera setup, but it is possible that some large birds could be detected at this distance.

To test the spatial accuracy of our calibration, we measured the distance between the tips of turbine blades and the tip of the hub in 23 frames chosen to represent a variety of spatial configurations. This resulted in a mean distance of 41.4 m, standard deviation of 1.05 m, and coefficient of variation of 0.02. Therefore, we can expect typical errors less than ± 1-2 m for this calibration setup. For comparison, a recently published study testing similar software for tracking birds and bats at wind turbines found errors up to ± 20 m (Matzner et al., 2020).

## 4. Discussion

### 4.1. Animal Tracking Applications

Videographic techniques are seeing expanded use for studies of wildlife (Cilulko, Janiszewski, Bogdaszewski, & Szczygielska, 2013; Christiansen et al., 2014; Gonzalez et al., 2016) ranging from animals in agricultural fields (Christiansen et al., 2014), to cetaceans (Seymour, Dale, Hammill, Halpin, & Johnston, 2017), bats and birds (Betke et al., 2008; Cullinan, Matzner, & Duberstein, 2015; Matzner, Cullinan, & Duberstein, 2015) and ants (Narendra & Ramirez-Esquivel, 2017). One major application of videography for studies of animals in the natural environment is counting populations. For example, 3-D thermal imaging has been used to show that bat colonies have only a small fraction of the number of individuals compared to earlier human counts (Betke et al., 2008). Thermal imaging has also been used in combination with radar and acoustics for monitoring migratory patterns of birds (Gauthreaux & Livingston, 2006; Horton, Shriver, & Buler, 2015).

A second major application of videography is studying animal movement patterns. Numerous studies have investigated the structure and rules underlying bird flocks (Ballerini et al., 2008; Evangelista et al., 2017) and schools of fish (Jolles et al., 2017). Videography has also been used for studying flight biomechanics of animals under natural conditions et al., 2015), and interactions between bats and birds and large structures such as wind turbines and oil and gas platforms (Horn et al., 2008; Cryan et al., 2014; Ronconi et al., 2015). Most of the studies described above have relied on custom software that is not widely available. The aim of the current study was to develop a robust, easy to use and free software package that could be used for these and other applications.

### 4.2 ThruTracker Capabilities

Here we present a new software package for 2-D and 3-D animal tracking. ThruTracker has several features not found in other freeware. These include easily adjustable procedures for 2-D and 3-D tracking; a tool for calibrating intrinsic parameters of thermal cameras; the ability to track and count hundreds of animals simultaneously; and the ability to make 3-D calibrations without dedicated calibration objects. We demonstrate these capabilities by counting bats leaving bridges and caves (Table 1; Figure 3) and making a 3-D calibration using only a wind turbine as a calibration object (Figure 4).

**Figure 4.**
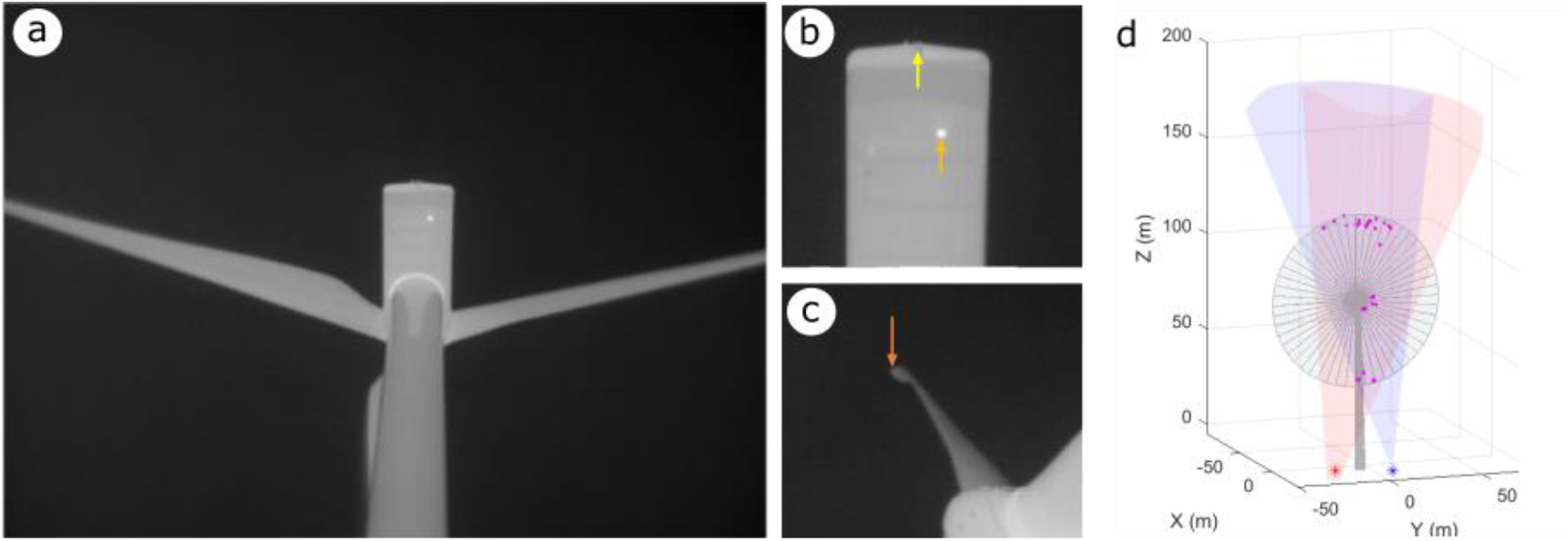
Extrinsic calibration of a wind turbine scene. (a) Example views from a single camera. (b, c) Examples of background points used for generating the calibration. These include a back corner of the nacelle (yellow arrow), a small hotspot of unknown origin (orange arrow) and a turbine tip (dark orange arrow). (d) Visualization of the calibrated scene. Views of two cameras are shown in light red and light blue shading. The wind turbine is shown as a grey outline but note that the turbine rotates about the vertical axis depending on wind direction. Magenta points show 3-D positions of points used for making the calibration.

The software is compatible with thermal and light-based imaging and most standard video formats (e.g., avi, wmv, mp4). ThruTracker uses an app-based environment with no coding required to make well-established detection and tracking algorithms widely available. Users simply import videos and select the detection and tracking options. These features make it easy to track moving animals under a variety of conditions.

### 4.3 Requirements and Limitations

ThruTracker uses a well-established background subtraction algorithm for object detection (Zivkovic, 2004; Zivkovic & Van Der Heijden, 2006). This method generates a rolling model of the background using a number of frames that can be specified by the user. This approach is best suited for stationary backgrounds and animals that are in near continuous motion. It has difficulty with animals that stay in one place; however, using more images for generating the background would help address this problem. ThruTracker aims to detect one point for each animal (the blob centroid). For detecting multiple body parts per animal one should consider deep-learning based approaches (Mathis et al., 2018; Pereira et al., 2019). ThruTracker has built-in compatibility for importing detections from other programs such as DeepLabCut (Mathis et al., 2018) for could be used for generating 3-D tracks from point clouds.

The main requirements for tracking in 3-D are 1) synchronized video are acquired from at least two fixed cameras 2) camera intrinsics are calibrated in the laboratory, and 3) some objects (including the focal animals themselves) are visible at a range of 2-D and 3-D positions within the calibrated volume. Without the requirements for dedicated calibration objects, it is now possible to calibrate nearly any volume in the lab or field. We demonstrate this workflow for generating 3-D calibrations at wind turbines, where it would be logistically challenging to put calibration objects in the airspace (Figure 4). Another approach would be to use the animals themselves as background points (Corcoran & Hedrick, 2019).

## 5. Conclusions

Technological development is driving price reductions and capability expansion in thermal and high-speed cameras, along with supporting equipment such as UAVs. However, the software required to make full use of these capabilities for research in fields as diverse as biomechanics, animal behavior, ecology, and population monitoring remains the province of specialized workflows in individual lab groups. ThuTracker provides an integrated, graphical, and user-friendly package to fill these needs, thus expanding the number of researchers able to make effective use of these emerging technologies.

## Acknowledgements

We thank Bethany Straw from the National Renewable Energy Laboratory for help with deploying thermal cameras. Iwona Kuczynska and Shelly Colatskie provided videos of gray bats exiting caves. William Valentine assisted with code development. Kacie Quigley, Madison Simmons and Jonas Håkansson conducted manual counts of bat exits from bridge and cave videos and Jonas Håkansson provided useful feedback on an earlier version of this manuscript.

## Authors’ Contributions

A.C. and T.H conceived of and developed the software. M.S. provided funding and guided software development. E.B. collected data used for testing and aided with data analysis. A.C. drafted the manuscript and all authors edited the manuscript and gave final approval for the publication.

## Data Availability Statement

ThruTracker installation files, a user manual and software tutorials are available at www.sonarjamming.com/thruTracker. Source code is available as a GitHub repository (https://github.com/AaronJCorcoran/ThruTracker).

